# Probing relaxed myosin states in hypertrophic cardiomyopathy by second harmonic-generation microscopy

**DOI:** 10.1101/2025.05.08.652664

**Authors:** Giulia Arecchi, Marica Dente, Weikang Ma, Beatrice Scellini, Nicoletta Piroddi, Marina Scardigli, Jingyuan Yu, Jing Zhao, Riccardo Cicchi, Ryo Kinegawa, Caroline Muellenbroich, Francesco Sera, Corrado Poggesi, Cecilia Ferrantini, Thomas C. Irving, Michael Regnier, Leonardo Sacconi, Chiara Tesi

**Affiliations:** Department of Experimental and Clinical Medicine, University of Florence, Florence, Italy; Illinois Institute of Technology, Chicago, IL, USA; College of Basic Medical Sciences, Dalian Medical University, Dalian, Liaoning, China; National Institute of Optics (INO) - CNR, Florence, Italy; School of Physics and Astronomy, University of Glasgow, Glasgow, United Kingdom; Department of Statistics, Computer Science and Applications "G. Parenti", University of Florence, Florence, Italy; School of Medicine & College of Engineering, University of Washington, Seattle, WA, USA; Institute of Clinical Physiology (IFC) - CNR, Florence, Italy

## Abstract

This study explores the use of polarized second-harmonic generation (pSHG) to investigate myosin conformation in the relaxed state, differentiating between the actin-available, disordered (ON) state and the energy-conserving, ordered (OFF) state. By shifting the ON/OFF equilibrium using both physical and chemical manipulations, we demonstrate the sensitivity of pSHG in quantifying the ON/OFF ratio in skeletal and cardiac tissues. Comparisons with X-ray diffraction measurements further validate our findings. Applying this approach to a sarcomeric mutation associated with hypertrophic cardiomyopathy, we show that R403Q/MYH7-mutated minipig ventricle tissue exhibits a higher ON fraction compared to controls. This difference is abolished under high concentrations of a myosin activator (2-deoxyATP) and an inhibitor (Mavacamten), indicating structural similarity between R403Q and controls in these two states. ATPase assays reveal increased resting ATPase activity in R403Q samples, which persists even in the presence of 2-deoxyATP, suggesting that the elevated energy consumption in the R403Q mutation is driven by both a population shift toward the ON state and enhanced myosin ATPase activity per motor head.

## INTRODUCTION

Exploring the molecular structural dynamics within living cells that drive biological functions poses significant challenges due to the wide range of spatial and temporal scales involved. A classic example is the collective chemomechanical action of myosin motors within the sarcomere, the fundamental structural unit of striated muscle. Contractile forces in muscle are generated through the "power stroke," a series of structural transitions in the actomyosin cross-bridge driven by ATP hydrolysis. Techniques such as cryo-electron microscopy (Cryo-EM) (1), electron paramagnetic resonance (EPR) (2), X-ray diffraction (3), fluorescence polarization (4), and, more recently, second-harmonic generation (SHG) (5) have provided detailed insights into the conformational changes of myosin motors during the power stroke.

Among these, SHG - a nonlinear optical process occurring in hyperpolarizable materials such as muscle - has proven particularly informative (6). In sarcomeres, SHG signals display a distinct pattern of alternating bright and dark bands that reflect the underlying striated architecture. This label-free method enables high-resolution probing of sarcomeres in intact tissues, making it an invaluable tool for studying sarcomere organization (7, 8). However, the potential of SHG extends beyond structural imaging. By varying the polarization of the incident light in single muscle fibres, it is possible to gain insights into structural states of actomyosin motors myosin conformations showing significant responsiveness to myosin conformational changes during muscle contraction (9, 10). Here, we further expand the application of SHG polarization anisotropy (pSHG) to investigate whether it can detect the distribution of conformations of myosin heads in relaxed striated muscle.

Emerging evidence suggests that, in relaxed muscle (i.e. when cytoplasmic calcium concentration falls below the threshold of thin filament activation) a fraction of myosin motors stick out of from the thick filament backbone while others are folded onto the thick filament surface. Folded heads assume a compact conformation or “interacting head motif structure” (11–15) shown by the reconstruction of the vertebrate thick filament of striated muscle. The fraction of heads folded onto thick filament has been correlated to a “sequestered” or “OFF” state that could be the structural counterpart of the functional "super-relaxed state" (18, 19) characterized by reduced (10 times) ATPase turnover compared to detached myosin heads in the "disordered relaxed " or “ON” state. Within sarcomeres, upon calcium activation, the fraction of myosin heads in the ON state participates in initial force production, while the remaining heads, folded on the filament, represent a “reserve” of energy-conserving motors. A thick filament regulatory mechanism has been proposed to release of myosin motors from a folded conformation, complementing the classic calcium-induced thin-filament mechanism (20), especially in the modulation of heart muscle contraction. Biochemical and structural studies have shown that the ratio between OFF and ON heads is significantly altered in congenital cardiovascular diseases such as hypertrophic cardiomyopathy (HCM), dilated cardiomyopathy (DCM), and heart failure (23, 24), leading to mechanical and energetic dysfunctions (25, 26). Understanding myosin conformational states in the relaxed state is therefore critical for elucidating their role in force generation and energy regulation.

In this study, we used pSHG to quantify imbalances in myosin head conformations in both skeletal and cardiac muscle, including cardiac samples carrying the R403Q-MYH7 mutation, a well-established cause of HCM (27, 28) already extensively characterized for its functional impact (29–31) also in association to the regulatory state of the thick filament (32). We examined how this mutation affects myosin conformation by modulating the ON-OFF balance either biophysically (e.g. varying temperature and substrate concentration) or pharmacologically (e.g. with activators 2-deoxyATP (33, 34) or inhibitors e.g., Mavacamten (35, 36). These findings were directly compared with complementary structural data obtained through X-ray diffraction. Additionally, ATPase activity assays were performed to correlate structural alterations with changes in energy consumption. By integrating structural and functional analyses of ON-OFF states, this work demonstrates a novel approach to elucidate the role of the R403Q mutation in the hypercontractility phenotype of HCM and to provide new insights into the energy-regulating mechanisms of myosin in cardiac muscle.

## MATERIALS AND METHODS

### Second-harmonic generation microscope and pSHG imaging

#### Experimental setup

A pulsed titanium-sapphire laser (Chameleon Ultra II, Coherent), operating at 740 nm with a pulse width of 140 fs and a repetition rate of 80 MHz, was used for excitation. A laser scanner (Rapid Multi Region Scanner, MBF Bioscience, Vidrio Technologies) equipped with two galvanometric mirrors and a resonant mirror was employed to scan the excitation beam across the sample. To vary the polarization of the excitation laser, we used a quarter-wave plate (WPQ10E-780, Thorlabs) to achieve a well-defined circular polarization, followed by a rotating polarizer (ELL14K and LPNIRB100-B, Thorlabs) to generate uniform linear polarization with limited intensity variation (I_PP_ < 1% over 2π). Polarized light was focused on the sample using a 40× immersion objective (W-Plan Apochromat, 40×, 1.0 NA; Zeiss), while the SHG signal was collected in the forward direction using a 40× water immersion objective (Achroplan, 40×, 0.8 NA; Zeiss). A narrow band-pass filter (390/10 nm; FBH390-10, Thorlabs) placed before a GaAsP photomultiplier module (H11706P-40; Hamamatsu) was used to selectively detect the SHG signal.

SHG images of muscle tissue were acquired without any exogenous labelling. To characterize the polarization dependence of the SHG signal (pSHG) in the muscle tissue, SHG images were collected with different polarization angles (φ) of the incident light relative to the fibre axis. The entire field of view (FOV) was first examined to assess sample quality and identify regions of interest (ROIs) where sarcomeres exhibited minimal structural disarray with a sarcomere length of 2.1 µm. Once well-organized regions were identified, ROIs of 14 × 14 µm were imaged at 50 frames per second while the polarization angle rotated at 45 rpm. The excitation laser power (measured at the sample) was set to 30 mW. For each SHG image, the SHG intensity was integrated over all pixels, producing a polarization anisotropy intensity profile (I_SHG_) as a function of the polarization angles (φ). Experimental data were then fitted with Equation 1, which describes the theoretical polarization modulation of I_SHG_ under the assumption of cylindrical symmetry (9):

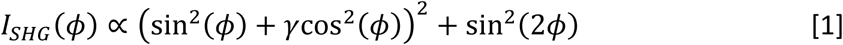

This fitting procedure allows extraction of the geometrical parameter γ, which is linked to the overall geometrical organization of SHG emitters within the excitation volume. The fitting procedure was carried out using a custom-made LabVIEW program (National Instruments, version 12.0f3 at 32 bit).

### Myofibril and skinned muscle strips

Rabbits (two males) were euthanized by pentobarbital administration (120 mg/kg) through the marginal ear vein, according to the procedure established by the European Union Council on the Use of Laboratory Animals (Directive 2010/63/EU), and using protocols approved by the Ethics Committee for Animal Experiments of the University of Florence. After careful dissection, the psoas muscle was cut into strips ∼0.5 cm wide, and samples were tied at rest length to rigid wood sticks, and stored at −20 °C for no more than 6 months in a 200 mM ionic strength Rigor solution (100 mM KCl, 2 mM MgCl_2_, 1 mM EGTA, 50 mM Tris, pH 7.0) supplemented with glycerol 50%. Single myofibrils or bundles of two to three myofibrils were prepared by homogenization of glycerinated psoas muscle as previously described (37).

Skinned muscle strips were obtained from rabbit psoas glycerinated muscle strips, from seven frozen minipig (males) hearts, and from seven mouse (males) hearts. We employed a knock-in minipig model carrying the R403Q mutation in MYH7, generated by introducing the mutation together with a BlastR selection cassette, which was subsequently excised via Cre-mediated recombination (36). The successful integration of the mutation was confirmed by Southern blot analysis using probes specific for MYH7 and BlastR, as well as by PCR with primers flanking the BlastR cassette, demonstrating its effective removal following Cre treatment. Isolated hearts from 1-month-old WT and R403Q male Yucatan pigs were provided by Exemplar Genetics Inc. Humane euthanasia and tissue collection procedures were approved by the Institutional Animal Care and Use Committees at Exemplar Genetics. The procedures were conducted according to the principles in the "Guide for the Care and Use of Laboratory Animals" (38), the Animal Welfare Act as amended, and with accepted American Veterinary Medical Association (AVMA) guidelines at the time of the experiments (39). Frozen left ventricular tissues were obtained from R403Q and WT minipig hearts and stored at -80 °C until the experiments. Each R403Q minipig was a sibling of a WT minipig.

C57BL/6J mice from Envigo (N=7, 6-7 months old) were used in this study. All animal handling and procedures were conducted in accordance with Directive 2010/63/EU of the European Parliament on the protection of animals used for scientific purposes. The experimental protocol was approved by the Italian Ministry of Health (reference code: 0DD9B.N.DZR). Gender issues have not been addressed in the present study. Mice were euthanized via inhaled isoflurane overdose (5%). Death was confirmed by the absence of a heartbeat, assessed through palpation, and the lack of thoracic movements. Once confirmed, the heart was explanted. Excised hearts were transported to the isolation station in a beaker containing 0.2 ml of heparin (5000 IU/ml) to prevent blood clotting, along with 10 ml of Krebs-Henseleit (KH) solution. Upon arrival at the isolation station, the hearts were immediately cannulated through the proximal aorta and retrogradely perfused with a KH solution modified to prevent spontaneous cardiac activity and to minimize contractures caused by shear damage during the dissection, and placed in a dissecting chamber under a binocular microscope. KH contained (mM): 120 NaCl, 5 KCl, 2 MgSO_4_, 1.2 NaH_2_PO_4_, 10 NaHCO_3_, 1.8 CaCl_2_ and 10 glucose, pH 7.38 equilibrated with HEPES buffer. During dissection, the potassium concentration was adjusted to 15 mM to halt spontaneous heartbeats. Additionally, 2,3-butanedione monoxime (BDM) at 20 mM was added to prevent contractions during surgery. The solution was continuously oxygenated with O₂.

The ventricular muscle strips were incubated overnight in a cold (4° C) standard relaxing solution added with 1% Triton X-100, to remove cellular membranes and all cytoplasmic organelles membranes, such as mitochondria and sarcoplasmic reticula. After this skinning procedure, the tissue was washed using a fresh relaxing solution containing protease inhibitors (AP). Under a stereoscope, the de-membranated strips were cut into smaller pieces, following the alignment of the fibres. Selected skinned muscle strips were attached to custom-made aluminium foil T clips and mounted horizontally between a force transducer and a length control motor in the experimental chamber. The length of the preparations was adjusted to a sarcomere length of ≅ 2.20 µm. The dimensions of the skinned strips were determined using an ocular micrometer mounted on the experimental microscope: on average, the preparations were 1 mm long, 140 µm wide, and 150 µm thick.

Myofibril and skinned preparations were pre-incubated in a series of relaxing solutions as follows: Relax (5 minutes), 2-deoxyATP (15 minutes), and Mavacamten (30 minutes), with imaging conducted after each incubation. Relax solutions: relaxing solutions were calculated as described previously at pH 7.0, and contained 10 mM of total EGTA, 5 mM MgATP, 1 mM free Mg^2+^, 10 mM 3-(N-morpholino) propane sulfonic acid, propionate, and sulfate to adjust the final solution to an ionic strength of 200 mM and a monovalent cation concentration of 155 mM. Phospho Creatine (10 mM) and Creatine Phospho Kinase (200 U ml^−1^) were added to all solutions to minimize alterations in the concentration of MgATP and its hydrolysis products. [Pi] resulting from contaminant inorganic phosphate was less than 150 µM. Different pCa > 8.00 with 10 mM EGTA were used for the experimental protocol:

- pCa > 8.00, 5 mM ATP

- pCa > 8.00, 5 mM 2-deoxyATP (Cat # C01577; Genscript, USA)

- pCa > 8.00, 5 mM ATP, and Mavacamten (Axon medchem BV -Groningen, The Netherlands; 50 µM in psoas muscle and 10 µM in cardiac preparation)

All solutions to which the samples were exposed contained a cocktail of protease inhibitors including leupeptin (10 µM), pepstatin (5 µM), phenylmethylsulphonyl fluoride (200 µM), and E-64 (10 µM), as well as NaN_3_ (500 µM) and 500 µM dithiothreitol.

For imaging in rigor a solution containing 100 mM KCl, 2 mM MgCl₂, 1 mM EGTA, and 50 mM Tris adjusted to pH 7.0 with HCl was used.

Additional experimental conditions were tested on rabbit psoas skinned preparations. Measurements were performed under conditions of lattice compression and at different temperatures.

For osmotic compression of the myofilament lattice, samples were measured in Relax solution supplemented with 5% (w/v) dextran (MW ∼500 kDa). Preparations were incubated for 15 minutes before imaging.

Temperature-dependent measurements were performed at 10°C, 15°C, and 20°C. The Relax solution was adjusted to the desired temperature, which was continuously monitored during acquisition using a microprobe placed in the experimental chamber. Preparations were incubated for 15 minutes at each temperature before imaging to allow thermal equilibration.

### Intact mouse cardiac preparations

Unbranched trabeculae extending from the free ventricular internal wall to the atrioventricular fibrous skeleton were dissected from mouse hearts by cutting their attachments: one end at the atrioventricular valve and the other at the ventricular wall. Trabeculae, with diameters ranging from 150 to 400 μm, were mounted between the hook-shaped extensor of two micromanipulators and placed in the imaging chamber equipped with two platinum electrodes for electrical stimulation. During imaging, the mouse trabeculae were stimulated at 0.1 Hz and continuously oxygenated. They were initially pre-incubated in KH solution for 15 minutes before the first imaging session. Subsequently, the preparations were exposed to KH containing 10 μM Mavacamten and incubated for 30 minutes prior to the second imaging session.

### ATPase measurements

#### Experimental setup

The experimental apparatus and the principle of the ATPase measurement were described in (40, 41). Briefly, the mounted muscle preparations were manually transferred between different baths, to expose them to different relaxing (pCa > 8.00) solutions: Relaxing, 2-deoxyATP Relaxing solution, and Mavacamten Relaxing solution, which were used subsequently for the measurements. The bath system consisted of an 80 μL well for incubating samples in relaxing solutions and a second well dedicated to the ATPase assay. This assay bath, with a volume of 30 μL, featured quartz windows to allow the transmission of near-UV light (340 nm) for measuring NADH absorbance. The bath was continuously stirred by motor-driven vibration of a membrane positioned at its bottom. The baths were milled in aluminium blocks (anodized) and mounted on top of an aluminium base, through which water was circulated to allow temperature control of all solutions (21 °C). ATPase activity measurements were normalized to the volume: length × cross-sectional area (CSA) of the muscle strip. The CSA was estimated in the setup assuming an elliptical shape: (width × depth × π)/4.

### Solutions

Different Relaxing solutions of pCa > 8.00 with 10 mM EGTA were used for the experimental protocol:

- Control Relaxing: pCa > 8.00, 6.2 mM ATP

- 2-deoxyATP Relaxing; pCa > 8.00, 6.2 mM 2-deoxyATP

- Mavacamten Relaxing; pCa > 8.00, 6.2 mM ATP, and Mavacamten 10 µM

All solutions contained: 100 mM BES (N,N-bis[2-hydroxyethyl]-2-aminoethane sulphonic acid); 6.63 mM MgCl. In addition, all solutions contained 0.25 mM NADH, 0.5 mg/mL pyruvate kinase (500 U/mg), 0.1 mg/mL LDH (912 U/mg), 10 μM oligomycin B. Oligomycine together with sodium azide 5 mM were added to suppress any residual mitochondrial ATPase or ATP synthase activity. The ionic strength of the solutions was maintained at 200 mM by adding the appropriate amount of potassium propionate (KProp). The pH was adjusted to 7.00 with KOH. Aliquots of these solutions (minus NADH) were stored frozen. On the day of the experiment, the solutions were thawed and the solutions were kept on ice until required.

All chemicals and enzymes, except for 2-deoxyATP and Mavacamten, were purchased from Sigma-Aldrich (Merck Life Science, St. Louis, MO, USA).

### Statistics

We used mixed-effects (multilevel) models to account for the potential non-independence of multiple measurements obtained from the same fibre or myocyte. Specifically, random-intercept models were used to estimate means and standard errors adjusted for the clustered structure of the data. In the mixed-effects models with random intercepts, fixed effects were also included to account for the experimental designs used in the different experiments (42, 43). Details of the statistical analyses performed for each measurement are provided in the corresponding figure legends. All analyses were performed using Stata version 19 (StataCorp LLC, College Station, TX, USA).

### X-ray diffraction measurements

#### Experimental setup

X-ray diffraction experiments were performed using the small angle X-ray diffraction instrument on the BioCAT beamline 18ID at the Advanced Photon Source, Argonne National Laboratory (44). The X-ray beam energy was set to 12KeV (0.1033 nm wavelength) at an incident flux of ∼5x10^12^ photons per second. The X-ray beam was focused to ∼250 μm × 250 μm (horizontal and vertical respectively) at the sample position and to ∼150 × 30 µm at the detector. The specimen to detector distance was about 3.5 m. The muscles were incubated in a customized chamber and experiments were performed at 28-30 °C. The muscles were stretched to a sarcomere length of 2.3 µm by monitoring the laser diffraction patterns from a helium-neon laser (633 nm). The X-ray patterns from either WT or R403Q samples were collected either in relaxing solutions with 100% ATP and then 100% 2-deoxyATP or in relaxing solution and then in relaxing solution with 50 μM Mavacamten on a MarCCD 165 detector (Rayonix Inc., Evanston IL) with a 1 s exposure time. The muscle samples were oscillated along their horizontal axes at a velocity of 1 - 2 mm/s to minimize radiation damage. The irradiated areas are moved vertically after each exposure to avoid overlapping X-ray exposures. 2-3 patterns were collected under each condition. The data were analysed using the MuscleX software package (45). The X-ray patterns were quadrant folded and background subtracted using the “Quadrant Folding” routine in MuscleX. The intensities and spacings of meridional and layer line reflections were measured by the “Projection Traces” routine in MuscleX as described previously (46). To compare the intensities under different conditions, and correct for any possible X-ray diffraction intensity changes due to radiation damage to the muscle, the measured intensities of X-ray reflections are normalized to the total diffuse background intensity as described previously (47). There were no significant changes in the radial width of the M3 reflection among different conditions, thus no correction for width was applied to the measured intensities. Values obtained from patterns under the same condition were averaged.

### Sample preparation and solutions

Previously frozen WT and R403Q left ventricular myocardium samples from 1-month old Yucatan mini-pig (male) were prepared as described previously (48, 49). Briefly, samples were defrosted in skinning solution (2.25 mM Na_2_ATP, 3.56 mM MgCl_2_, 7 mM EGTA, 15 mM sodium Phospho Creatine, 91.2 mM potassium propionate, 20 mM Imidazole, 0.165 mM CaCl_2_, Creatine Phospho Kinase 15 U/ml with 15 mM 2,3-Butanedione 2-monoxime (BDM) and 1% Triton-X100 and at pH 7) at room temperature for 30 min before splitting to smaller bundles (500 µm in diameter and variable lengths). The fibre bundles were further skinned in fresh skinning solution at room temperature for 2-3h. Muscles were then washed with fresh relaxing solution (2.25 mM Na_2_ATP, 3.56 mM MgCl_2_, 7 mM EGTA, 15 mM sodium Phospho Creatine, 91.2 mM potassium propionate, 20 mM Imidazole, 0.165 mM CaCl_2_ Creatine Phospho Kinase 15 U/ml at pH 7) for ≍10 minutes, repeated 3 times. The tissue was dissected into ≍200-300 μm diameter, 3-5 mm long fibre bundles and clipped with aluminium T-clips. The preparations were then stored in cold (4°C) relaxing solution with 3% dextran for the day’s experiments. The muscles were incubated in a customized chamber, and experiments were performed at 28-30 °C.

## RESULTS

### pSHG sensitivity to myosin conformation in rabbit psoas skeletal muscle

pSHG microscopy (Fig. 1A) was first applied to demembranated rabbit psoas skeletal muscle, which has a well-organized structure, to investigate whether it can detect myosin conformations in relaxed muscle. SHG images are characterized by a remarkable alternation of light and dark bands, reflecting the sarcomeric striations typical of the muscular tissue (Fig. 1B). pSHG recordings were performed in selected regions of interest (ROIs) in demembranated fibres in rigor and in relaxing solution in the absence or presence of a high concentration of the myosin motor inhibitor Mavacamten or the myosin activator 2-deoxyATP, a naturally occurring ATP analogue. The four experimental conditions highlight different pSHG intensity modulations, which can be well described by the theoretical polarization modulation of I_SHG_ (Eq. 1), resulting in distinct γ values (Fig. 1C). Repeated measurements on different rigor fibres yielded an average value of *γ_Rigor_* = 0.63 ± 0.02 (mean ± SEM) while the same measurement performed on relaxed fibres produced an average value of *γ_Relax_* = 0.39 ± 0.01 (Fig. 1D). The large statistically significant difference between *γ*_Rigor_ and *γ*_Relax_ confirms the capability of pSHG to probe distinct conformation changes of myosin motors. Notably, by replacing 5 mM ATP with 5 mM 2-deoxyATP or adding 50 µM Mavacamten to the relaxed (5 mM ATP) state, we observed a significant variation in the *γ* value relative to baseline condition. The positive variation with 2-deoxyATP was *γ_dATP_* = 0.49 ± 0.02 and a negative variation with Mava was *γ_Mava_* = 0.28 ± 0.01.

**Figure 1:**
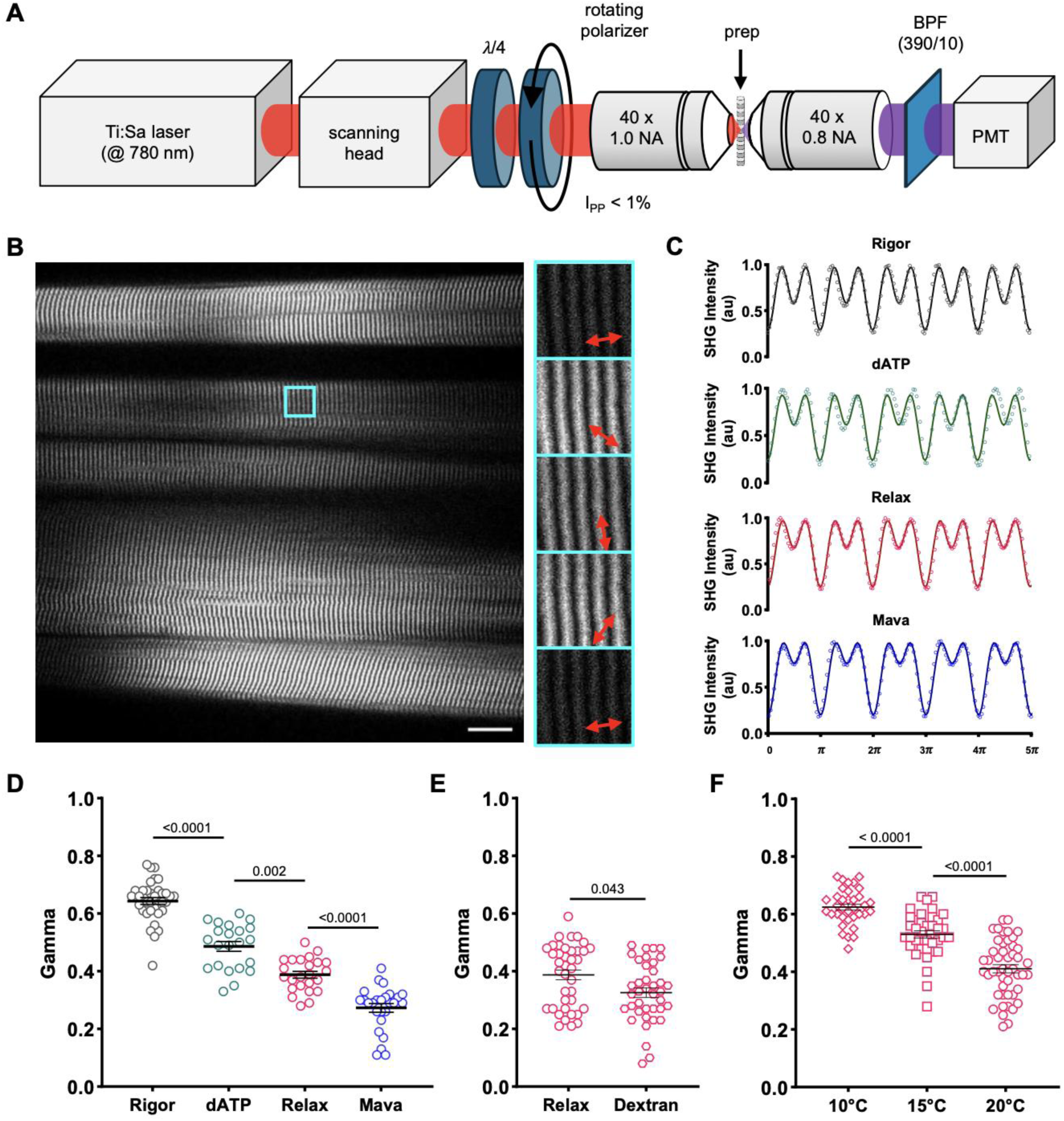
pSHG sensitivity on myosin conformation in psoas skeletal muscle. A) Optical scheme of the SHG microscopy setup. The excitation beam’s optical path is shown in red, while the emission path is in purple. Abbreviations: λ/4, quarter-wave plate; BPF, bandpass filter; PMT, photomultiplier. B) SHG image of a demembranated rabbit psoas fibre. Bright bands correspond to sarcomeric A bands. Scale bar: 20 µm. The cyan square highlights a representative region of interest (ROI) selected for acquisition (13 × 13 µm). The right column shows Selected ROI illuminated at different polarization angles. C) Representative SHG polarization curves in rigor, after the exposure to a high concentration of 2-deoxyATP (dATP, 100%), in relaxation, and after exposure to a high concentration of Mavacamten (Mava; 50 µM). Circles represent raw data, and the continuous lines indicate the best fit of Eq. 1, yielding a best-fit parameter of γ_Rigor_ = 0.68 in rigor, γ_dATP_ = 0.53 with dATP, γ_Relax_ = 0.36 in relax, and γ_Mava_ = 0.27 with Mavacamten. D) Graph illustrating γ values among the Rigor state, after exposure of 2-deoxyATP (dATP), Relax state, and after exposure of Mavacamten (Mava) in rabbit skinned psoas fibres. Data were obtained from 9 psoas fibres, with multiple ROIs collected across each strip, producing 35 data points for the Rigor state, 22 for dATP, 43 for relaxation, and 50 for Mavacamten. Data are reported as mean ± S.E.M. On comparing Rigor vs. dATP, Relax, and Mava we included conditions as fixed effects, while on pairwise comparisons between dATP, Relax, and Mava, the strip was also added as a fixed effect. E) Graph illustrating γ values in relaxing solution and in relaxing solution supplemented with 5% dextran in rabbit skinned psoas fibres. Data were obtained from 3 psoas fibres, with multiple ROIs collected across each strip, producing 30 data points for Relax state and 30 data points for dextran. On comparing Relax vs. 5% dextran we included condition as a fixed effect along with strip. F) Graph illustrating γ values in relaxing conditions at three different temperatures (10 °C, 15 °C, and 20 °C) in rabbit skinned psoas fibres. Data were obtained from 5 psoas fibres, with multiple ROIs collected across each strip, producing 35, 41, and 48 data points at 10 °C, 15 °C, and 20 °C, respectively. On comparing the effect of temperature we included temperature as a fixed effect along with strip. For each comparison, we reported the p-value associated with the Wald test of the specific coefficient. In each experiment, we adopted the Bonferroni correction to evaluate the significance of the p-values.

The reduction of *γ* observed in the presence of Mavacamten is consistent with a conformational change in myosin as it approaches the thick filament backbone (35). Conversely, 2-deoxyATP binds to the myosin catalytic site and promotes a shift toward the ON state, resulting in an increase in the γ value. Of note, the increase in the γ value observed with 2-deoxyATP is not linked to actomyosin complex formation, as no detectable increase in passive force was observed in the presence of 2-deoxyATP (see Fig. S1). A similar behaviour was observed in isolated myofibrils, where γ values were comparable to those measured in skinned preparations (see Fig. S2), further supporting the efficacy of our technique in probing myosin conformational states.

To further explore the sensitivity of pSHG, we manipulated myosin conformation by changing lattice spacing and temperature under relaxing condition. We found that, in the presence of dextran (Fig. 1E), the γ value slightly decreased, consistent with a shift of myosin toward the thick filament backbone, as previously reported (50, 51). Finally, lowering the temperature (Fig. 1F) caused a significant increase in γ, indicating a shift toward the ON state, as previously described using x-ray diffraction (51).

### pSHG sensitivity in demembranated and intact cardiac preparations

We further evaluated the pSHG sensitivity to distinct relaxed myosin states by applying the same experimental approach to mouse ventricular preparations (Fig. 2). The cardiac SHG images also displayed regular alternation of bright and dark bands, visually underscoring the uniformity of the sarcomere architecture. We made measurements of skinned mouse cardiac samples in rigor, relaxed, 100% 2-deoxyATP and 10 µM Mavacamten. Consistent with our observations in the psoas muscle, we observed a significant difference of γ value between rigor (*γ_rigor_* = 0.71 ± 0.01) and relaxed (*γ_Relax_* = 0.31 ± 0.01) and, under relaxing condition, the γ value increased after the addition of 2-deoxyATP (*γ_dATP_* = 0.47 ± 0.02) and decreased after the addition of Mavacamten (*γ_dATP_* = 0.24 ± 0.01; Fig. 2B).

**Figure 2:**
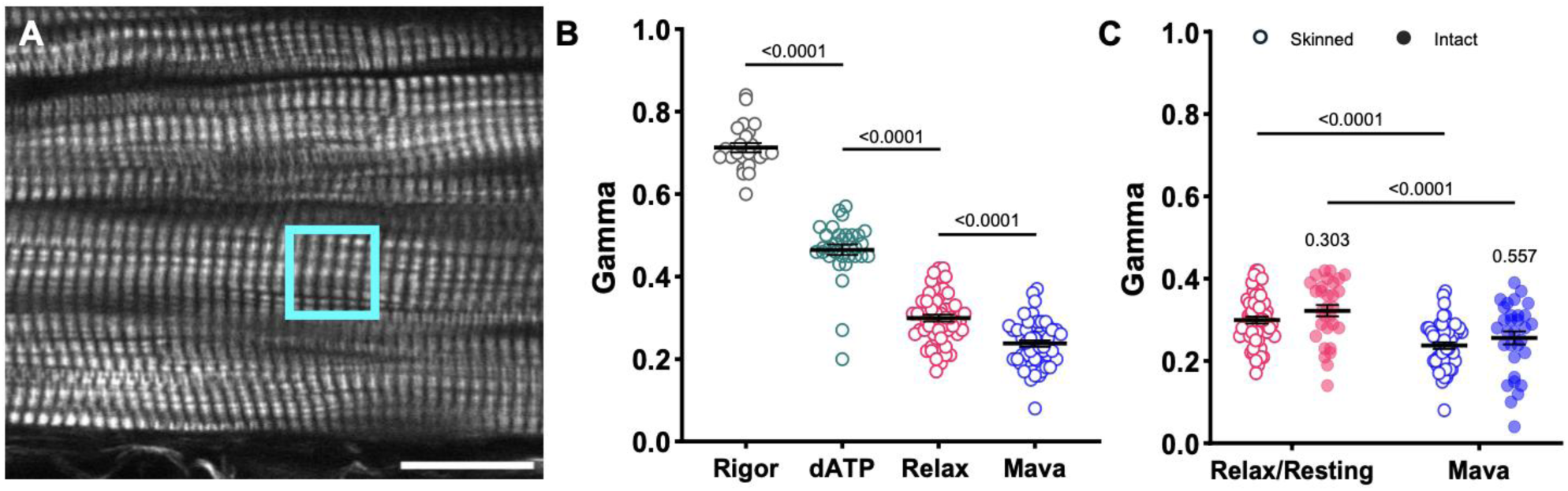
pSHG sensitivity on demembranated and intact cardiac preparations. A) Representative SHG image of a demembranated mouse ventricular wall strip. Borders between individual cells are visible. Scale bar: 20 µm. The cyan square highlights a representative region of interest (ROI) selected for acquisition (13 × 13 µm). B) Graph illustrating γ values among the Rigor state, after exposure to 2-deoxyATP (dATP), Relax state, and after exposure to Mavacamten (Mava) in mouse skinned cardiac samples. Data were obtained from 14 skinned cardiac samples dissected from 5 mice. Multiple ROIs were collected across each strip, producing 24 data points for the Rigor state, 32 for dATP, 86 for relaxation, and 52 for Mavacamten. Data are reported as mean ± S.E.M. On comparing Rigor vs. dATP, Relax, and Mava we included conditions as fixed effects, while on pairwise comparisons between dATP, Relax, and Mava, the strip was also added as a fixed effect. C) Comparison of γ values in mouse demembranated versus intact cardiac preparations. The graph shows γ values measured in both skinned (open circles) and intact (filled circles) multicellular cardiac preparations. Data were collected first in a Relax solution and subsequently after the addition of 10 µM Mavacamten. For intact preparations, stimulation was applied at a frequency of 0.1 Hz, with imaging performed during the long diastolic phase. Data were obtained from 5 skinned cardiac samples dissected from 3 mice and 6 intact trabeculae dissected from 3 mice. Multiple ROIs were collected across each strip, producing 55 data points for the Relax state and 52 for Mavacamten in skinned preparation while 30 data points for the resting state and 30 for Mavacamten were obtained from intact preparations. Data are reported as mean ± S.E.M. On comparing Relax/Resting vs. Mava, a stratified analysis was performed in the two preparation groups (Skinned and Intact). We included condition as a fixed effect, along with strip. To compare the two preparations in the two conditions (Relax/Resting vs. Mava), we considered a mixed-effect model with preparation as a fixed effect. For each comparison, we reported the p-value associated with the Wald test of the specific coefficient. In each experiment, we adopted the Bonferroni correction to evaluate the significance of the p-values.

We also examined whether γ values were comparable in skinned and intact cardiac preparations. To this end, we conducted pSHG measurements in intact murine trabeculae stimulated at 0.1 Hz, with a diastolic phase duration compatible with the pSHG scan. The γ values in intact samples were consistent with those observed in demembranated strips in the relaxing solution and varied to a similar extent after pharmacological treatment with 10µM Mava (Fig. 2C). These data demonstrated the effectiveness of pSHG in probing distinct structural conformations of myosin *in vivo* at rest, reinforcing the robustness and reliability of the method across different experimental conditions.

### ON-OFF myosin state analysis by pSHG in R403Q-MYH7 mutation

Driven by these results, we employed pSHG to probe the ratio between the disordered-relaxed (ON) and sequestered (OFF) states in a minipig model carrying an HCM-associated mutation (R403Q-MYH7). Myocardial disarray is one of the structural abnormalities classically associated with HCM, and its presence contributes to contractile dysfunction and energetic impairment (52). To minimize polarization artefacts potentially caused by structural disarray - particularly evident in pathological preparations (Fig. 3A), the entire sample was initially examined to carefully identify ROIs where sarcomeres displayed minimal disarray.

**Figure 3:**
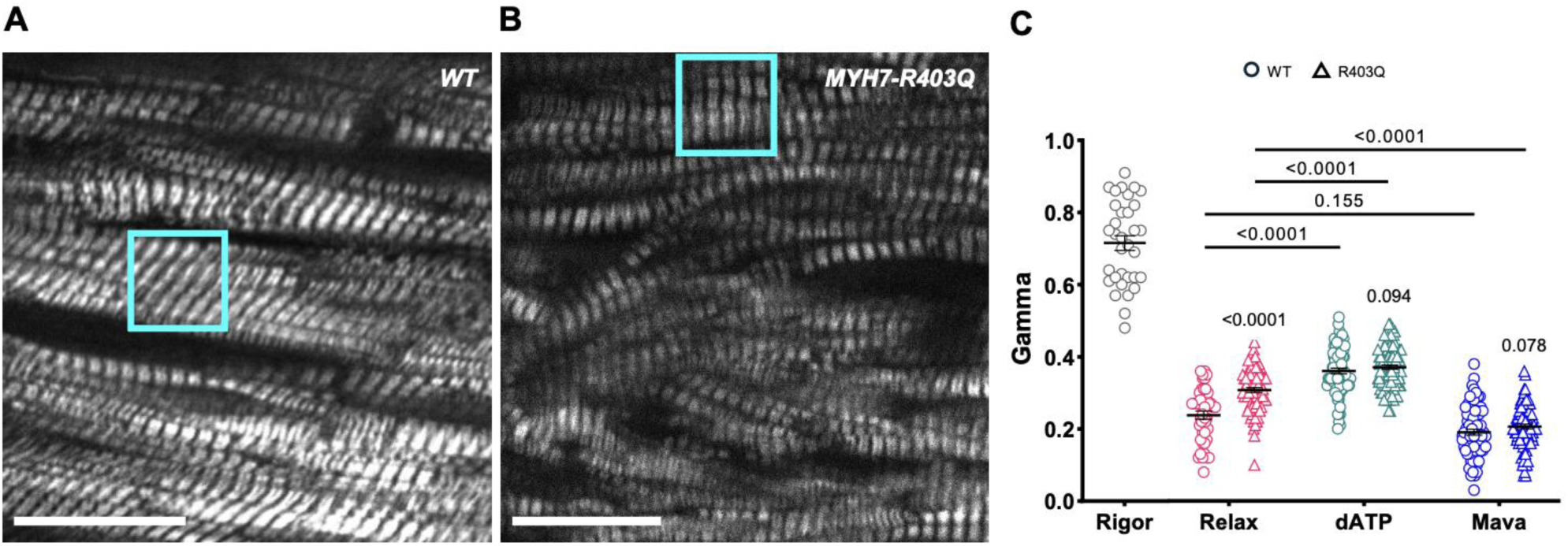
pSHG in R403Q-MYH7 mutation. A) SHG image of a sample from a wild-type (WT) minipig, and B) sample from a minipig carrying the MYH7-R403Q hypertrophic mutation. Cyan squares highlight representative regions of interest (ROIs) selected for acquisition (13 × 13 µm). A marked difference is evident between healthy and pathological tissue, with the mutated tissue exhibiting a high degree of structural disarray. Scale bars: 20 µm. C) Pharmacological manipulation of skinned preparations from WT and R403Q minipigs. The graph shows γ values obtained under different conditions: in Rigor, in Relax solution, after exposure to 100% 2-deoxyATP (dATP), and following treatment with 10 µM Mavacamten (Mava). Data were obtained from 11 skinned cardiac samples dissected from 4 WT minipig and 11 skinned cardiac samples dissected from 3 R403Q-MYH7 minipig. Multiple ROIs were collected across each strip, producing 33 data points for the Rigor state in WT samples only. For WT and R403Q-MYH7 samples, respectively, 45 and 76 data points were obtained for the Relax state, 77 and 103 for 2-deoxyATP, and 76 and 91 for Mavacamten. Data are reported as mean ± S.E.M. On comparing Rigor vs. Relax, dATP, and Mava we included conditions as fixed effects. Pairwise comparisons between dATP, Relax, and Mava were performed on each group (WT and R403Q) using mixed-effect models with condition and strip as fixed effects. The comparison between genotypes on each condition (dATP, Relax and Mava) was performed using mixed-effect models with genotype as fixed effects. The likelihood ratio test was performed to test the interaction between genotype and condition. For each comparison, we reported the p-value associated with the Wald test of the specific coefficient. In each experiment, we adopted the Bonferroni correction to evaluate the significance of the p-values.

Data acquisition was collected in these well-organized regions. From pSHG measurements in the relaxed condition (Fig. 3B), we observed a γ value in R403Q-MYH7 that was significantly higher than that measured in sibling wild-type minipigs (WT). Notably, the higher γ value in R403Q-MYH7 is consistent with an increased level of ON state myosin heads associated with this mutation.

We next performed pharmacological manipulations to compare the effects of 2-deoxyATP (promoting a shift toward the ON state) and Mavacamten (driving the equilibrium toward the OFF state) in preparations expressing the R403Q mutation and WT. As expected, the application of 100% 2-deoxyATP or 10 µm Mavacamten resulted in an increase and a decrease in γ values, respectively, in both preparations. Intriguingly, in the presence of either molecule, the original differences between the WT and R403Q groups disappeared, suggesting a similar myosin structural conformation between WT and R403Q under these conditions. Consistent with our findings in psoas and mouse ventricular preparations, all values were significantly lower than the γ value observed in rigor (Fig. 3C).

To enhance the robustness of our findings, complementary X-ray diffraction measurements were performed (Fig. 4). X-ray diffraction patterns obtained from R403Q minipig model in a relaxing solution show a marked decrease in the intensity of the myosin-based M3 meridional reflection compared to the WT littermates. Among myosin-based reflections, the M3 meridional reflection originates from the axial repeat of myosin motors, indicating an increase in disordered heads in R403Q. Consistently, the spacing of the M6 meridional reflection (SM6), originating from a periodic mass distribution in the thick filament backbone, is enhanced in R403Q compared to WT. As expected, upon the application of 2-deoxyATP and Mavacamten on the same strips (paired measurements), we observed that R403Q preparations, compared to WT, there was reduced effect of 2-deoxyATP but larger effect of Mavacamten. These findings further support that the R403Q mutation shifts myosin towards the ON state, underlying the hypercontractility characteristic of R403Q-induced HCM.

**Figure 4:**
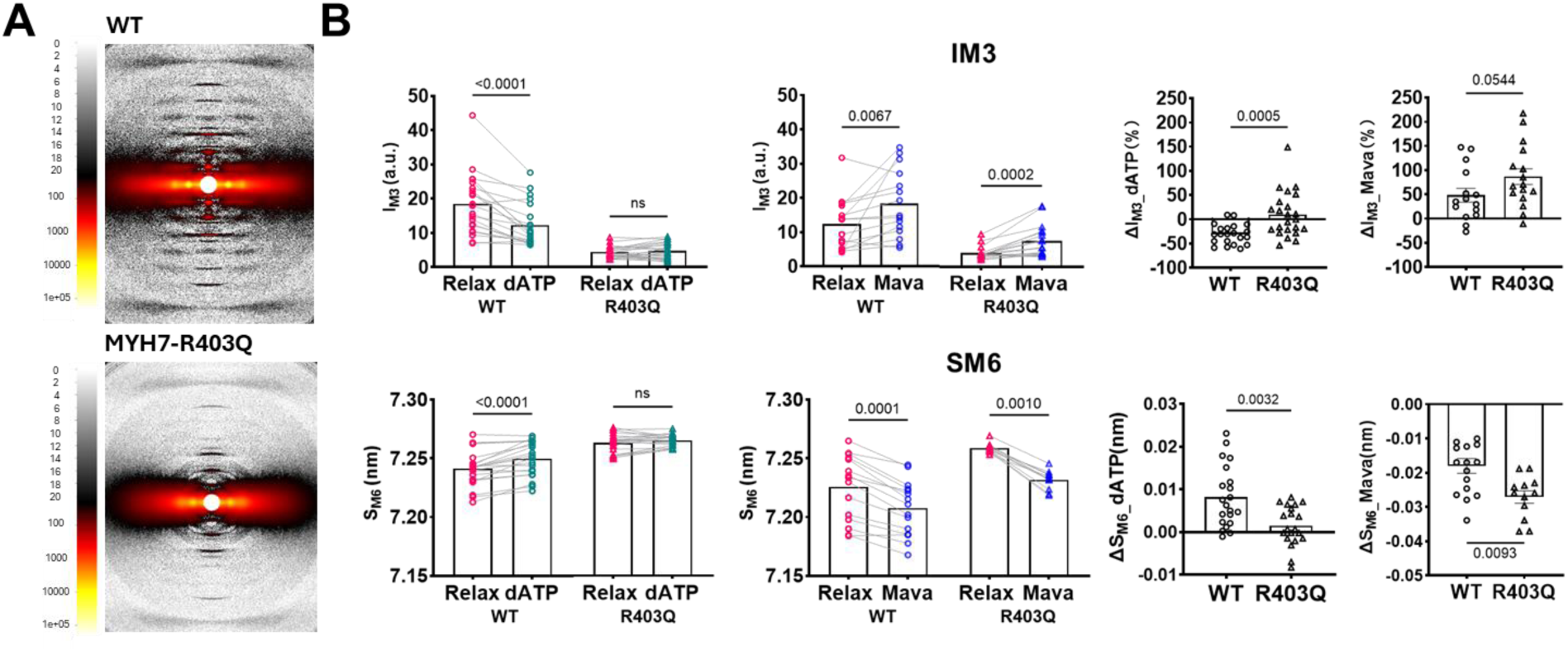
X-ray diffraction in R403Q-MYH7 mutation. A) Representative X-ray diffraction patterns from wild-type (WT) and R403Q minipig ventricular strips in relaxing solution. B) Top row: Graphs comparing changes in M3 reflection intensity (IM3) between WT and R403Q samples under different conditions. The first two graphs show intensity differences between Relaxed and 2-deoxyATP (dATP) treated samples (WT and R403Q, respectively) and between Relaxed and Mavacamten (Mava) treated samples. The third and fourth graphs illustrate ΔIM3, comparing the changes in IM3 between WT and R403Q samples with 2-deoxyATP and Mavacamten treatments. Bottom row: Graphs comparing the spacing of the M6 meridional reflection (SM6) between WT and R403Q samples under different conditions. The first two graphs display shifts in SM6 between Relaxed and 2-deoxyATP treated samples and between Relaxed and Mavacamten treated samples for both groups. The third and fourth graphs show ΔSM6, comparing shifts in SM6 between WT and R403Q with 2-deoxyATP and Mavacamten treatments. Data were obtained from > 15 skinned cardiac samples dissected from 2 WT minipig and 12 skinned cardiac samples dissected from 2 R403Q-MYH7 minipigs.

### ATPase activity measurements in R403Q mutant myocardium

To provide a more comprehensive understanding of the implications of the R403Q mutation, alongside the pSHG analysis, we performed ATPase activity measurements to evaluate the specific effects of the R403Q mutation on sarcomere energetics at rest. This correlative approach enabled us to link structural observations with functional assessments, deepening our understanding of the R403Q mutation’s influence on the ON and OFF states of myosin.

As expected, a significant difference in ATPase activity was observed between WT and R403Q preparations, with the latter exhibiting markedly higher energy consumption compared to the WT counterparts (Fig. 5). This finding aligns with the structural analysis, where mutant preparations showed a shift toward the ON state. Exposure to a high concentration of Mavacamten (10 µM) resulted in a global reduction in energy consumption in both preparations, bringing ATPase activity to comparable levels. These results are consistent with the structural analysis, which showed a complete shift of myosin heads toward the OFF state in both WT and R403Q samples. On the other hand, in the presence of 2-deoxyATP, which by itself is associated with increased ATPase activity (53), a significant disparity in energy consumption between the two groups remained, with the R403Q mutation associated with a higher energy requirement. Under the same conditions, the pSHG data indicated that both WT and R403Q samples underwent comparable structural conformation.

**Figure 5:**
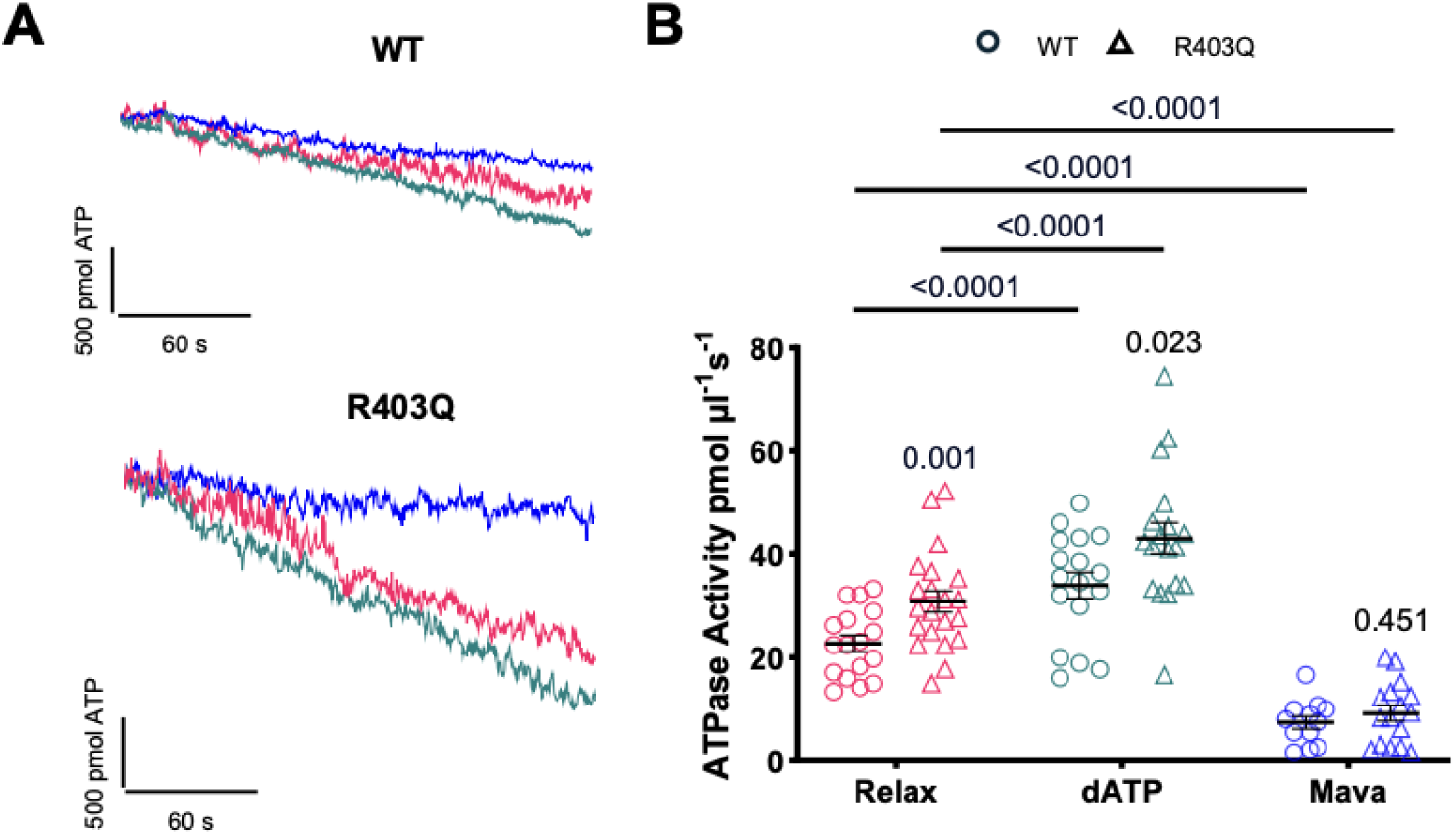
ATPase activity in R403Q mutation. A) Representative traces of NADH absorbance from wild-type (WT) minipig preparations (left column) and preparations from minipigs carrying the MYH7-R403Q hypertrophic mutation (right column) under different conditions: Relax, 100% 2-deoxyATP (dATP), and 10 µM Mavacamten (Mava). B) Graph illustrating ATP consumption across different sample conditions: in Relaxing solution, after exposure to 100% 2-deoxyATP, and following treatment with 10 µM Mavacamten. Data were obtained from 17 skinned cardiac samples dissected from 3 WT minipig and 21 skinned cardiac samples dissected from 3 R403Q-MYH7 minipig. Each preparation is associated with a data point in the three conditions. Data are reported as mean ± S.E.M. Pairwise comparisons between Relax, dATP, and Mava were performed on each group (WT and R403Q) using mixed-effect models with condition and strip as fixed effects. The comparison between genotypes on each condition (Relax, dATP, and Mava) was performed using mixed-effect models with genotype as fixed effects. The likelihood ratio test was performed to test the interaction between genotype and condition. For each comparison, we reported the p-value associated with the Wald test of the specific coefficient. In each experiment, we adopted the Bonferroni correction to evaluate the significance of the p-values.

This observation suggests that the increased energy consumption associated with the R403Q mutation is not solely explained by a differences in the balance between the structural ON and OFF states in WT and R403Q myocardium (53).

## DISCUSSION

Thanks to the coherent summation of hyper-Rayleigh scattering from the amide groups (HN-CO) of polypeptide chains - which act as second-harmonic emitters - SHG signals can be endogenously detected in certain biological structures that possess the necessary spatial organization and symmetry. The organization of myosin into helical filaments and their arrangement into cylindrically symmetric, repetitive structures along the fibre provide an ideal configuration for generating SHG (9). In this context, the distribution of second-harmonic emitters can be probed by analysing the polarization modulation of SHG intensity (pSHG), which enables the derivation of a γ value associated with the emitters distribution angle relative to the principal axis of symmetry (fibre axis). However, to accurately correlate the γ value with the protein’s structural conformation, it is essential to understand how the second-harmonic emitters are organized within the protein under investigation. By reconstructing each contributing second-harmonic emitter within the actomyosin motor array at the atomic scale, Nucciotti et al. (5) established a direct relationship between myosin conformation and the γ value. Notably, this revealed an overall trend: the γ value increases as the overall angle of myosin relative to the thick filament backbone becomes larger and decreases as this angle is reduced. Although this study shows that it is not possible to achieve a unique myosin structural assignment from γ, it may nonetheless provide a useful means to distinguish between ‘sequestered’ (OFF state) and ‘disordered’ (ON state) myosin molecules in the relaxed state.

In this study, we employed pSHG to investigate the structural conformation of myosin in the relaxed state in skeletal (rabbit psoas) and cardiac (mouse and minipig) muscle by means of the γ value. Under relaxing conditions, pSHG analysis revealed a γ_Relax_ value of 0.39 ± 0.01 (mean ± SEM) in psoas skeletal muscle, 0.31 ± 0.01 in mouse cardiac muscle, and 0.22 ± 0.02 in minipig cardiac muscle. The difference in γ values between skeletal and cardiac muscle can be related to distinct structural arrangements of myosin heads in the relaxed thick filament, as well as variations in the basal ON/OFF state ratio. pSHG measurements under conditions that maximise OFF-state contributions (in the presence of high doses of Mavacamten) showed an overall decrease in γ values, whereas measurements under conditions that minimize OFF-state contributions (in the presence of 100% 2-deoxyATP) showed an overall increase in γ values. Interestingly, even under pharmacological conditions that shift myosin heads toward saturation of ON or OFF states, significant differences in γ values persisted between skeletal and cardiac muscle (Fig. S3). In skeletal muscle, particularly in the psoas, the predominant myosin heavy chain (MyHC) isoforms are IIx (MyHC1) and IIb (MyHC4), which are characteristic of type II muscle fibres, also known as fast-twitch fibres. The dominant isoform in mouse cardiac ventricular muscle is alpha myosin heavy chain (MyHC6 or α-MyHC), typical of cardiac tissue in small mammals and atrial tissue in larger mammals (56, 57). Conversely, in minipig ventricular cardiac muscle, the main isoform is beta myosin heavy chain (MyHC7 or β-MyHC), characteristic of ventricular muscle in larger mammals, including humans (58, 59). Various myosin isoforms polymerize to confer distinct structural organizations to the thick filament backbone and distinct profiles of heterogeneity of the myosin heads “crowns” along the structure possibly leading to differences in γ values here observed, independent of, and additive to, the contribution from the ON/OFF state ratio of the myosin heads (16, 17).

Notably, the impact of Mavacamten and 2-deoxyATP in shifting the γ value differs between skeletal and cardiac muscle, suggesting a difference in ON/OFF state fraction across muscle types. In detail, assuming that these two pharmacological manipulations maximize the OFF and ON states, respectively, we estimate ON/OFF ratios at room temperature of 0.50 in psoas skeletal muscle, 0.30 in mouse cardiac muscle, and 0.20 in minipig cardiac preparations. The estimated ON/OFF ratios are consistent with values obtained using X-ray diffraction and fluorescence-based methods under comparable experimental conditions in both skeletal (51) and cardiac muscle (60, 61). In contrast, the ON/OFF state ratio found here does not align with prior estimates obtained through mantATP biochemical assays (18, 36, 62–64). This discrepancy suggests that an overestimation of fast cycling heads (ON state) may occur even after correction for non-specific binding of the nucleotide. This contributes to the ongoing discussion in the context of the numerous unknown factors involved in correlating the structural and functional states of the myosin motor (65–67).

pSHG was also employed in preparations carrying the R403Q mutation, showing a clear increase in γ values in the R403Q mutation, consistent with the hypothesis of an increased ON population compared to WT. X-ray diffraction patterns obtained from the same R403Q minipig model show a marked decrease in the intensity of the third-order meridional reflection (M3) in the R403Q preparations compared to the WT littermates. In addition, the spacing of the sixth-order meridional reflection (SM6) increases significantly in the R403Q minipig preparations. The increase in the thick filament backbone periodicity, indicated by the increased SM6, and a reduction in the degree of helical ordering of the myosin heads, indicated by the reduced M3 intensity, are characteristic signatures of the structurally-defined OFF-ON transition of myosin (68). In line with this observation, we found that R403Q is less sensitive to 2-deoxyATP compared to WT. This is likely due to R403Q already increasing the ON population, such that the 2-deoxyATP effect on the ON/OFF population is diminished. Our additional observation of a pronounced effect of Mavacamten increase M3 intensity and SM6 spacing provides evidence that the R403Q mutation alters the conformation of myosin in the thick filament, favouring the ON state (32).

Taken together, these results support the widely accepted hypothesis that many HCM-associated β-cardiac myosin mutations destabilize the structural OFF states of thick filament-assembled myosin. This occurs by disrupting key interactions within the motor head and/or neck, leading to an increase in the number of heads available for interaction with actin and altering the ON/OFF ratio of motors participating in the cross-bridge cycle (26, 29, 69–74). This view is consistent with the reported increase in resting energy consumption in the presence of R403Q β-cardiac myosin in cardiac sarcomeres. An increase in the resting energy consumption associated with the R403Q mutation is also confirmed by our ATPase measurements, where we observe a higher ATP turnover in R403Q minipig cardiac strips compared to WT. The functional effects of the R403Q mutation on myosin motor activity have been reported to vary depending on the experimental system and assay employed, with studies showing both reduced and enhanced motor performance (75–77). We previously investigated the relationship between cross-bridge kinetics and energetics in human cardiac tissue by comparing samples from R403Q patients with mutation-negative HCM controls (31). Basal ATPase activity was also elevated in R403Q human samples compared with mutation-negative HCM controls (31), further supporting the hypothesis that the mutation alters energetic balance in the relaxed state.

Additionally, in the presence of a high concentration of Mavacamten, ATPase measurements are significantly reduced in both R403Q and WT, resulting in statistically indistinguishable values. These data are in full agreement with the coupled pSHG measurements, which indicate that in the presence of Mavacamten, R403Q and WT are represented with a similar ON/OFF state fraction.

On the other hand, it is important to note that ATPase measurements carried out under 2-deoxyATP conditions show an additional significant increase in energy consumption for both WT and R403Q groups. It has been reported that 2-deoxyATP increases ATPase of acto-HMM (53) and myosin S1 (66), demonstrating the presence of increased basal ATPase activity independent of the ON/OFF configuration of myosin structure. Here we show that increased hydrolytic activity of myosin with 2-deoxyATP is also present despite changes in myosin structure induced by R403Q mutation. Data obtained from our correlative approach, which combines parallel structural and functional investigations, suggest that the R403Q mutation not only destabilizes the structural OFF states of thick filament-assembled myosin but also associates with an increase of intrinsic activity of myosin motors .

The capability of pSHG to probe myosin dynamics structurally in physiological contexts, represents a novel complementary approach to well-established atomic-scale techniques like X-ray diffraction. This study also demonstrated the ability to monitor myosin conformational changes under relaxing conditions also *in vivo*, where pSHG showed γ values that were not statistically different from those observed in the same muscle in the skinned configuration. This finding not only opens the possibility of applying this method to intact multicellular preparations but also reinforces the reliability of the geometrical parameter γ as a tool for selectively probing myosin states.

The micron-scale resolution of SHG microscopy inherently limits a comprehensive assessment of molecular conformation, as well as the fine spatial characterization of the zonal distribution of OFF/ON states along the thick filament. In particular, it does not allow resolution of the structural heterogeneity associated with the presence of MyBP-C, as described in single-molecule studies (32, 78) and interference X-ray diffraction analyses (79), nor of the effects related to the phosphorylation state of myosin light chains (80).

Notwithstanding these limitations, the ability to probe the myosin state in vivo through polarization anisotropy without the need for labelling, combined with its relatively modest technological requirements, makes pSHG an accessible and valuable tool for microscopy facilities, particularly when compared with atomic-scale techniques such as X-ray diffraction.

### Future perspectives

Based on these findings, we propose that this technique could serve as a valuable tool for gaining deeper insights into the physiology and pathophysiology of muscle. For example, studying how temperature and lattice factors affect the ON/OFF ratio estimated by pSHG (see Fig. 1E,F as example), in conjunction with ATPase measurement, could help to answer the still open question of how the structural IHM state correlates with the OFF or "SRX" state in resting muscle (67). In addition, the sensitivity of this technique to the ON/OFF ratio could be further explored across species by employing different concentrations of Mavacamten and 2-deoxyATP. Finally, the possibility of probing the ON/OFF ratio within the sarcomere using a high-numerical-aperture objective to achieve sub-sarcomere resolution opens the opportunity to explore the heterogeneity of thick filament structure (81) in a label-free modality.

In pathological contexts, numerous applications and studies could be envisioned. For example, the ON/OFF ratio at rest is thought to be primarily regulated by cardiac protein C (82), highlighting mutations in this protein - especially in cases of haploinsufficiency - as a critical area of research.

Additionally, future studies might explore mutations in thin filaments, which appear to elicit comparable effects (83). This suggests that a variety of mutations may converge on a shared molecular ’phenotype.’

Moreover, the temporal resolution of our pSHG approach (seconds) could enable the observation of myosin in stationary states, such as during the plateau phase of isometric contraction. To capture the rapid dynamics of the myosin working stroke, which occurs on a millisecond timescale, the temporal resolution could be further enhanced by limiting pSHG measurements to two orthogonal angles, one parallel and one perpendicular to the fiber. This implementation could significantly improve the time resolution, expanding the applicability of this technique to the study of working stroke dynamics.

Lastly, the lab-scale setup of pSHG is well-suited for further implementation in high-throughput screening systems, where the ON/OFF ratio could be assessed across different pathological settings and patients in a multi-well configuration (84).

## ACKNOWLEDGMENTS

We thank Annibale Biggeri for preliminary statistical analysis not included in the present work, and Massimo Reconditi for fruitful discussions. This research project has been supported by the European Union’s Horizon 2020 programme under grant agreement No. 952166 (REPAIR) and HORIZON-HLTH-2023-TOOL-05 SMASH-HCM grant agreement number No.101137115; by the Italian Ministry of University and Research (MUR) under grant PRIN2022 #2022F3NENH; by the European Union’s NextGenerationEU Programme with the I-PHOQS Infrastructure; by National Heart Lung and Blood Institute Grant (R01HL171657, WM), National Institute of Health; by the MUR - Dipartimenti di Eccellenza 2023-2027 to the Department of Experimental and Clinical Medicine of the University of Florence; by University of Florence funding - Large Equipment 2021. This research used resources of the Advanced Photon Source, a U.S. Department of Energy (DOE) Office of Science User Facility operated for the DOE Office of Science by Argonne National Laboratory under Contract No. DE-AC02-06CH11357. This project is supported by grant P30 GM138395 from the National Institute of General Medical Sciences of the National Institutes of Health, P30 AR074990 from the National Institute of Arthritis and Musculoskeletal and Skin Diseases, and NIH R01 HL128368 from the National Heart, Lung, and Blood Institute. The content is solely the authors’ responsibility and does not necessarily reflect the official views of the National Institute of General Medical Sciences or the National Institutes of Health.

