## Supplementary Figures for "Probing relaxed myosin states in hypertrophic cardiomyopathy by second harmonic-generation microscopy"

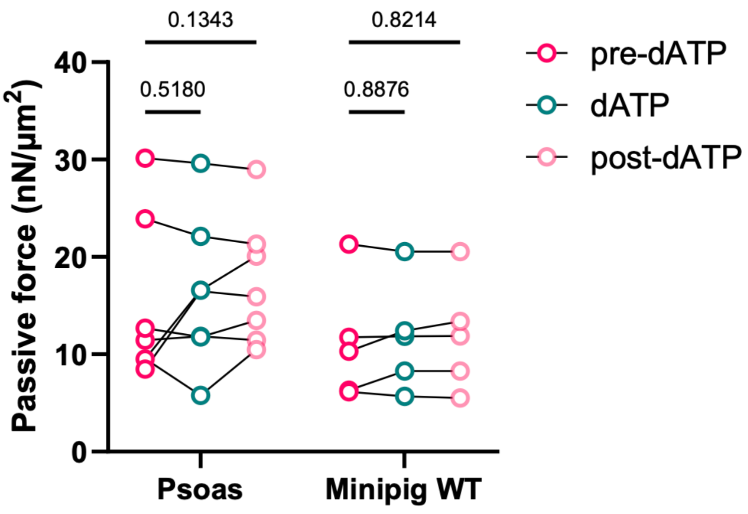


**Figure S1. Passive force measurements in psoas and minipig myofibrils.** The graph shows passive force recorded in myofibrils initially exposed to an ATP-containing solution (pre-dATP), followed by exposure to a 100% 2-deoxyATP solution (dATP), and subsequently returned to the ATP-containing solution (post-dATP), in both psoas myofibrils (average sarcomere length: 2.77 ± 0.07 μm) and minipig myofibrils (average sarcomere length: 2.20 ± 0.08 μm). Statistical analysis was performed using two-way ANOVA followed by Tukey's post hoc test.

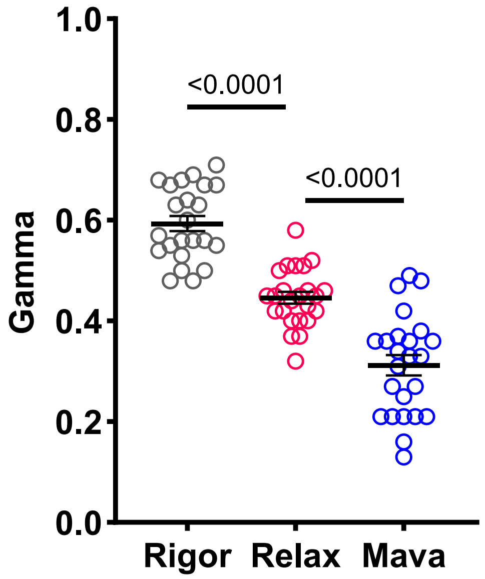


**Figure S2: Rabbit psoas myofibrils.** Graph illustrating γ values among the Rigor state, Relax state, and after exposure of Mavacamten (Mava), in rabbit skinned psoas myofibrils. Data are reported as mean ± S.E.M. A one-way repeated measures ANOVA was performed with a Tukey post-hoc correction.

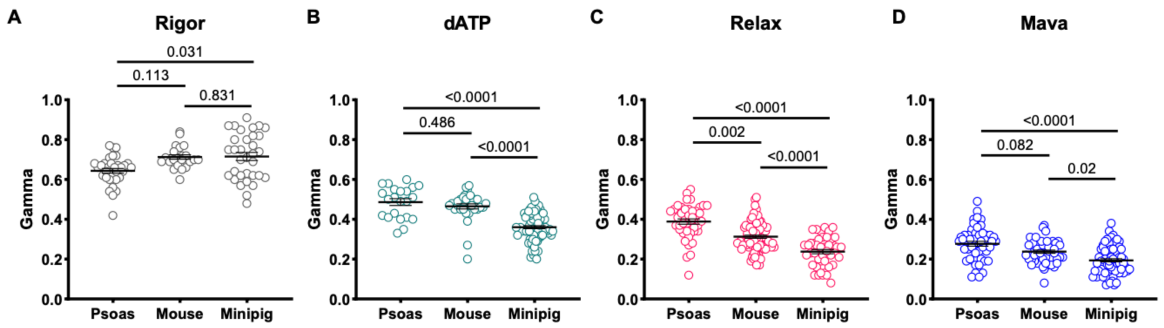


**Figure S3. Comparison of γ values among different species under distinct chemically modulated conditions.** Comparison of γ values measured in skinned preparations from rabbit psoas muscle, mouse cardiac muscle, and minipig cardiac muscle under the following conditions: Rigor solution (A), Relax solution with 100% 2-deoxy-ATP (dATP) (B), Relax solution (C), and Relax solution after treatment with high concentrations of Mavacamten (50 µM in psoas, 10 µM in mouse and minipig preparations) (Mava) (D). Data are presented as mean ± S.E.M. For each condition (Rigor, dATP, Relax and Mava), the species effect was evaluated considering mixed models with species as fixed effects.
